# Wnt5a and Notum Influence the Temporal Dynamics of Cartilaginous Mesenchymal Condensations in Developing Trachea

**DOI:** 10.1101/2024.09.02.610014

**Authors:** Natalia Bottasso-Arias, Megha Mohanakrishnan, Sarah Trovillion, Kaulini Burra, Nicholas X. Russell, Yixin Wu, Yan Xu, Debora Sinner

## Abstract

The trachea is essential for proper airflow to the lungs for gas exchange. Frequent congenital tracheal malformations affect the cartilage, causing the collapse of the central airway during the respiratory cycle. We have shown that Notum, a Wnt ligand de-acylase that attenuates the canonical branch of the Wnt signaling pathway, is necessary for cartilaginous mesenchymal condensations. In Notum deficient tracheas, chondrogenesis is delayed, and the tracheal lumen is narrowed. It is unknown if Notum attenuates non-canonical Wnt signaling. We observed premature tracheal chondrogenesis after mesenchymal deletion of the non-canonical Wnt5a ligand. We hypothesize that Notum and Wnt5a are required to mediate the timely formation of mesenchymal condensations, giving rise to the tracheal cartilage. Ex vivo culture of tracheal tissue shows that chemical inhibition of the Wnt non-canonical pathway promotes earlier condensations, while Notum inhibition presents delayed condensations. Furthermore, non-canonical Wnt induction prevents the formation of cartilaginous mesenchymal condensations. On the other hand, cell-cell interactions among chondroblasts increase in the absence of mesenchymal Wnt5a. By performing an unbiased analysis of the gene expression in Wnt5a and Notum deficient tracheas, we detect that by E11.5, mRNA of genes essential for chondrogenesis and extracellular matrix formation are upregulated in Wnt5a mutants. The expression profile supports the premature and delayed chondrogenesis observed in Wnt5a and Notum deficient tracheas, respectively. We conclude that Notum and Wnt5a are necessary for proper tracheal cartilage patterning by coordinating timely chondrogenesis. Thus, these studies shed light on molecular mechanisms underlying congenital anomalies of the trachea.

## INTRODUCTION

Patterning the developing respiratory tract requires precise interactions between the endoderm-derived epithelium and the surrounding mesenchyme (Caldeira et al., 2021; Nasr et al., 2020; Shannon and Hyatt, 2004). This feedback is essential to determine the branching pattern of the developing lung (Zhang et al., 2022), pulmonary epithelial cell differentiation (Brownfield et al., 2022), as well as cell differentiation of the mesenchyme and epithelium of the central airways of the respiratory tract (Bottasso-Arias et al., 2022; Zhou et al., 2022). Our previous studies have identified Notum as a target of epithelial Wls-mediated signaling that influences mesenchymal condensation and cartilaginous ring morphology. Furthermore, Notum is an essential component of a negative feedback loop attenuating Wnt/β-catenin signaling in developing mesenchyme (Gerhardt et al., 2018).

Wnt5a, a Wnt ligand triggering non-canonical response in developing respiratory tract, plays roles in epithelial cell differentiation and alveolarization, partially by inhibiting Wnt/β-catenin signaling via mechanisms dependent on Ror1/2 (Baarsma et al., 2013; Li et al., 2020). Aberrant expression of Wnt5a is also associated with cancerous lung disease, pulmonary fibrosis, bronchopulmonary dysplasia (BPD), and hyperoxia injury (Sucre et al., 2020). Non-canonical signaling exerts a critical role in the convergent extension movements of cells via planar cell polarity (PCP), supporting the elongation of different structures during development (Tada and Heisenberg, 2012), the position and branching pattern in developing lung (Zhang et al., 2022), tracheal length (Kishimoto et al., 2018), and the orientation and assembly of the epithelium and smooth muscle cells of the trachealis muscle (Kishimoto et al., 2018; Russell et al., 2023b). Notably, in developing trachea, we detected a precise pattern of expression wherein Notum expression is positioned between epithelial Wnt7b (a ligand inducing Wnt/β-catenin) and mesenchymal-produced non-canonical Wnt5a. However, whether Notum can modulate non-canonical Wnt signaling and its effect on morphogenesis and disease outcome remains unclear.

Previous studies have shown that anomalous cartilage patterning leads to congenital malformations such as tracheomalacia, a genetic structural abnormality of the trachea with weak cartilage lining, and tracheal stenosis, with both impairing breathing. We identified compound heterozygous single nucleotide variant (SNV) in ROR2 in patients diagnosed with complete tracheal ring deformity (CTRD), a condition characterized by tracheal stenosis (Sinner et al., 2019). ROR2 encodes the receptor of WNT5A supporting a role for the ligand Wnt5a via ROR2 mediated signaling in the patterning of the central airways of the respiratory tract in humans.

Chondrogenesis leading to mature cartilage is a tightly regulated process encompassing morphogenetic events and phenotypic changes (Goldring et al., 2006). The first step in chondrogenesis is cell condensation and the formation of condensed cell aggregates. The process of mesenchymal condensation primarily involves an increase in local cell density, mediated by local cell movements without altered proliferation (Klumpers et al., 2014). These local cell rearrangements are mediated by passive extracellular matrix ECM-driven movements and by dragging and pushing by neighboring cells (Oster et al., 1985) (Cui et al., 2005). Studies by Barna and Niswander (Barna and Niswander, 2007) demonstrated in an *in vitro* model with single cell imaging resolution, that the process of mesenchymal condensation is very dynamic requiring sorting, migration and cell shape changes partially mediated by Sox9 and Bmp4. At this early stage in mesenchymal condensation formation, the major components of the ECM are hyaluronic acid and fibronectin (FN) (Knudson and Toole, 1985; Kulyk et al., 1989). Simultaneously, cell–cell contacts mediated through N-cadherin and neural cell adhesion molecule (N-CAM), are increased to facilitate cell–cell communication likely triggering the onset of chondrogenic differentiation (Widelitz et al., 1993). This model agrees with the notion that a high cell density is required for chondrogenesis (Mauck et al., 2002). Supporting this concept, studies on tooth development demonstrated that cell compaction causes the activation of a genetic cascade necessary for specific cell induction (Mammoto et al., 2011). While cellular events driving the cell condensation have been unveiled, less is known about how genetic clues are integrated and transduced into forces mediating cell compaction.

In the present work, we sought to investigate the role of Notum influencing the Wnt5a-mediated signaling in tracheal mesenchymal condensations that give rise to the tracheal rings. We hypothesize that Notum represses Wnt5a-mediated non-canonical Wnt signaling, which is required for tracheal mesenchyme patterning and cartilage formation. We determined that Wnt5a, similarly to Notum, is necessary for the timely condensation of chondroblasts, and its ablation causes precocious condensation affecting cartilaginous ring morphology and number. In vivo, Notum fine-tunes the Wnt canonical and non-canonical levels in developing tracheal mesenchyme to promote the formation and patterning of tracheal cartilage.

## MATERIAL AND METHODS

### Mouse breeding and genotyping

Animals were housed in a pathogen-free environment and handled according to the protocols approved by CCHMC Institutional Animal Care and Use Committee (Cincinnati, OH, USA). Generation of Notum^150/150^ mice was previously described (Gerhardt et al., 2018). Adult Notum mice were kept in heterozygosis. Wnt5a ^f/f^ (Jackson lab # 026626) were mated with Dermo1 ^Cre/wt^ (Jackson lab# 008712) to generate Dermo1 ^Cre/wt^; Wnt5a ^f/wt^ mice and rebred to Wnt5a ^f/f^ generating embryos of genotype Dermo1Cre; Wnt5a^f/f^ mice (Ryu et al., 2013). Notum^300/150^ mice were bred with Wnt5a^f/f^ (Jackson lab # 026626) mice to generate Notum^300/150^ Wnt5a^f/w^, and the resulting mouse rebred to Wnt5a^f/f^ to generate Notum^300/150^ Wnt5a^f/f^. Notum^300/150^ mice were bred with Dermo1^cre/wt-^ (Jackson lab# 008712) to generate Notum^300/150^ Dermo1^cre/wt-^. The Notum^300/150^ Dermo1^cre/wt-^ were then crossed with a Notum^300/150^ Wnt5a^f/f^ to produce Notum^150/150^ Wn5a^f/wt-^Dermo1^cre/wt-^ embryos. Sox9KIeGFP mice was previously described (Chan et al., 2011) (Jackson laboratories # 030137). Genotypes of transgenic mice were determined by PCR using genomic DNA isolated from mouse-tails or embryonic tissue. γSMA mouse was previously described (Bottasso-Arias et al., 2022; Russell et al., 2023b). Primers utilized for genotyping are provided as Supplementary material. (Supplementary table 1).

### DNA constructs

Plasmid utilized for transfection were pGL2 Top Flash, (Sinner et al, 2004), pcDNA3 Full Length Notum, (GenScript), pGL2ATF2 (a gift from Dr. Niehrs)(Ohkawara and Niehrs, 2011), pGL3AP1(Addgene), pcDNA6 activeWnt5a, and pcDNA6 active Wnt3a (Addgene).

### Transfections and luciferase assay

NIH3T3 cells were cultured in 48 well plate, at 37° C and 5% CO_2_. Cells at 60%-70% confluency were transfected with a total of 50 ng of plasmidic DNA using Fugene (Promega) according to manufacturer’s instructions. These reporters were co-transfected with Wnt3a, Wnt5a and/or Notum expression plasmids (Gerhardt et al., 2018). Cells were harvested 24 hours post transfection, washed with PBS, and lysed using a passive lysis buffer reagent (Promega). Luciferase activity was determined over 10 s integration time using a luminometer (Promega).

### Dissociation and FACS of embryonic trachea

Cell dissociation was performed following a previously published protocol (Bottasso-Arias et al., 2022). Per each sample, five E13.5 *γSMA^eGFP^* tracheas/litter were pooled together and dissected in cold PBS and dissociated to single cells using TrypLE Express (phenol-red free, Thermo, 12604013) at 37°C for 10 minutes, followed by trituration for 30 seconds at RT. Cells were washed twice with FACS buffer (1mM EDTA, 2% FBS, 25mM HEPES in phenol-red free HBSS). To identify epithelial cells, cells were stained with APC anti-mouse CD326/EpCAM (Invitrogen, ref 17-5791-82, used at 1:50) at 4°C for 30 minutes followed by two washes with FACS buffer. Cells were resuspended in FACS buffer and passed through a 35 µm cell strainer. To stain dead cells, Sytox Blue nucleic acid stain (Thermo, S11348, used at 1 µM) was added to the cell suspension. Cells were sorted using a BD FACS Aria I and II. Single live “chondroblast enriched cell population” was collected after size selection and gating for Sytox-negative, EpCAM-negative, and eGFP negative cells. Cells were sorted directly into RNA lysis buffer (Zymo Research Quick RNA Micro kit, S1050) for isolation of RNA. Cell isolation strategy was confirmed by qRT-PCR (Supplement Figure1) and as previously shown (Bottasso-Arias et al., 2022; Russell et al., 2023a). Data is shown for three independent samples.

### Transcriptomic analyses

RNA-sequencing data were generated and compared from E11.5 *Dermo1Cre; Wnt5a ^f/f^* tracheas vs E11.5 Wnt5af/f tracheas and E13.5 *Notum^150/150^* vs E13.5 Notum ^300/300^ chondroblasts (GEO repository under GSE260707). Differentially expressed genes were identified using Deseq2 (E11.5 Wnt5a tracheas N=5 Controls, N=4 Mutants; Notum E13.5 sorted chondroblasts N=3 controls, N=4 mutants) (Anders & Huber, 2010; Lawrence et al., 2013). Fragments Per Kilobase (FPKM) values were calculated using Cufflinks (Trapnell et al., 2010). Differentially expressed genes were identified with the cutoff of a p-value <.05, FC>1.5 and FPKM>1 in over half of the replicates in at least one condition. The same approach was utilized for gene expression analysis of E13.5 control chondroblast, epithelium and smooth muscle cells isolated by FACS. (GEO repository GSE241175). Heatmaps were generated using normalized counts generated by DEseq2 and pheatmap or from RNA-seq fold changes. Functional enrichment was performed using Toppfun and hits relevant to this project were visualized in a - log10 (pvalue) bubble chart. System models were created using IPA’s Path Designer.

### Whole mount staining

Tracheal lung tissues isolated at E11.5-E14.5 were subject to whole mount immunofluorescence as previously described (Sinner et al., 2019). Embryonic tissue was fixed in 4% PFA overnight and then stored in 100% Methanol (MeOH) at -20°C. For staining, wholemounts were permeabilized in Dent’s Bleach (4:1:1 MeOH: DMSO: 30%H2O2) for 2 hours, then taken from 100% MeOH to 100% PBS through a series of washes. Following washes, wholemounts were blocked in a 2% BSA (w/v) blocking solution for two hours and then incubated, overnight, at 4°C in primary antibody diluted accordingly in the blocking solution. After five one-hour washes in PBS, wholemounts were incubated with a secondary antibody at a dilution of 1:500 overnight at 4°C. Samples were then washed three times in 1X PBS, transferred to 100 % methanol, through a series of washes in dilutions of methanol, and cleared in benzyl-alcohol benzyl-benzoate (Murray’s Clear). Images of wholemounts were obtained using confocal microscopy (Nikon A1R). Imaris imaging software was used to convert z-stack image slices obtained using confocal microscopy to 3D renderings of wholemount samples.

### Immunofluorescence staining and quantification

Embryonic tissue was fixed in 4% PFA overnight and embedded in paraffin or OCT to generate 7μm sections. For general immunofluorescence staining, antigen retrieval was performed using 10mM Citrate buffer, pH6. Slides were blocked for 2 hours in 1XTBS with 10% Normal Donkey serum and 1% BSA, followed by overnight incubation at 4°C in the primary antibody, diluted accordingly in blocking solution. Slides were washed in 1X TBS-Tween20 and incubated in secondary antibody at 1:200, in blocking solution, at room temperature for one hour, washed in 1X TBS-Tween20, and cover-slipped using Vecta shield mounting media with or without DAPI. Fluorescent staining was visualized and photographed using automated fluorescence microscopes (Nikon). Area of Sox9 positive cells was determined using Image J. Intensity of the N-cadherin stainings were quantified using NIS elements. Antibodies utilized in this manuscript have been previously validated by our laboratory and other investigators. Source, references, and dilution of primary and secondary antibodies used have been provided as Supplementary material (Supplementary table 2).

### Embryonic tracheal-lung culture

Sox9KI (Sox9-GFP) Embryonic tracheas were harvested at E12.5 and cultured in air-liquid interphase for 42 hrs as described (Hyatt et al., 2002). Images were obtained at frequent intervals and timelapse videos were recorded overnight between 24 and 42 hours employing an EVOS M700 microscope (Supplementary videos 1-4). Samples were treated with vehicle (DMSO), ABC99 (25uM Sigma SML2410), KN93 (5μM Millipore Sigma 422708) or JNK inhibitor II (8μM Millipore Sigma 420119). Wnt5a conditioned media was obtained from L cells (ATCC cat no. CRL-2647CRL-2814) and was diluted 50% with growth media (DMEM 5% FBS) for the experiment.

### Tracheal mesenchymal cell Isolation and culture

Primary cells were isolated as previously described (Bottasso-Arias et al., 2022; Gerhardt et al., 2018). Briefly, E13.5 tracheas of at least five embryos of the same genotype were isolated, washed in 1X PBS, dissociated in TrypLE express (Gibco) and incubated for 10 minutes at 37°C. After incubation, tissue was pipetted until cell suspension formed. Cells were seeded in flasks containing MEF tissue culture media composed of DMEM (ATCC 30-2002), 1% penicillin/streptomycin (Gibco 15140122), 2% antibiotic/antimycotic (Gibco 15240062), and 20% non-heat inactivated FBS (R&D S11150). Only mesenchymal cells were attached, as we confirmed expression of Sox9, Col2a1 and Myh11 but no expression of Nkx2.1 was detected (Bottasso-Arias et al., 2022).

### Wound-healing assay

Procedure was performed according to the manufacturer’s instructions using Culture-Inserts 2 Well (Ibidi) on a 24-well plate. A suspension of 8×10^5^ trachea MEF cells per mL isolated from control and *Wnt5a* and *Notum* knockout cells were plated on each side of the insert. A day after seeding, the inserts were removed, and a live cell nuclear dye (NucBlue ReadyProbe R10477, Invitrogen) was added to the cell media. The 24-well plate was then placed in an Evos M7000 microscope with an incubator (Thermo Fisher) (37°C and 5% CO2) overnight and timelapse images were taken with 20-minute time intervals between images for brightfield and DAPI channels. The total incubation time was 24 hours. ImageJ was used to measure the wound area from the brightfield images at various time points.

### Directional migration

Haptotaxis assay was performed using the CytoSelect 24-well kit (8um, fibronectin coated, colorimetric format) according to the manufacturer’s instructions (CBA-100- FN, Cell Biolabs Inc.). Briefly, 300 μL of a suspension of 1.0 x 10^6^ cells/ml was seeded inside the insert for each cell type (Notum control and mutant cells, Wnt5a control and mutant cells). Cells were allowed to migrate to the bottom of the insert coated with fibronectin for 24 hours. Afterwards, the remainder of the cells was removed from the inside of the insert and the cells that migrated to the bottom of the insert coated with fibronectin were stained with the dye. After washes and drying the inserts, the dye was solubilized and 100uL were transferred to a 96-well plate to measure absorbance at 560nm in a plate reader (Spectra MR Dynex Technologies). N=5 for each genotype.

### Cell adhesion

Cell adhesion assays were performed using micromasses plated with the different cell types (Notum control and Notum mutant, Wnt5a control and Wnt5a mutant). 50,000 cells per micromass were seeded in a 24-well plate. After allowing the micromasses to adhere for 2 hours, they were washed with PBS and fixed with 100% methanol. The micromasses were then stained with 0.1% Crystal Violet diluted in distilled water. The staining was removed, and the wells were washed with distilled water. A lysis-resuspension solution consisting of 10% methanol and 5% glacial acetic acid was added to each well, and cells were resuspended. 60 µL of the cells were transferred to a 96-well plate and the absorbance was read between 570-585 nm. A comparison between the absorbance of control and mutant cells was used relatively to determine cell adhesion.

### In-Situ Hybridization

The procedure was performed according to a protocol developed by Advanced Cell Diagnostics (ACD) (Wang et al., 2012). In situ probes were designed by ACD. Slides were baked and deparaffinized. In situ probes were added to the slides and hybridization was performed for 2 hours at 40°C, followed by several rounds of amplification steps. For fluorescence detection, opal dyes were utilized to detect the localization of the transcripts. After mounting with permanent mounting media (ProLong Gold, Thermo), slides were photographed using a wide field Nikon fluorescent microscope.

### Statistics

Quantitative data were presented as mean ± standard error. For animal experiments, a minimum of three different litters for each genotype were studied. Experiments were repeated at least twice with a minimum of three biological replicates for each group. Statistical analysis was performed using Graph Pad Prism ver.10 for MacOS. Statistically significant differences were determined by paired T-test, or one-way or two-way ANOVA repeated measures followed by post hoc pairwise multiple comparison procedures (Dunnet or Holm-Sidak test). Significance was set at P<0.05.

## RESULTS

### Notum attenuates non-canonical Wnt signaling, and is needed for proper cartilage development

Published studies have demonstrated that Notum and Wnt5a are involved in forming and patterning the tracheal cartilaginous rings. At E13.5, Notum is localized in the ventral subepithelial mesenchyme of the trachea, and Wnt5a is localized in the ventral mesenchyme of the trachea, just outside the localization of Notum. In both cases, Wnt5a and Notum were found to be in the region where tracheal cartilage forms and wherein Col2a1 and Sox9 are expressed (Fig 1b). Notably, Wnt7b (a ligand inducing Wnt/β-catenin dependent signal) is observed in the epithelium overlapping with Nkx2.1 and Cdh1 (Fig 1a,b). The complementary and distinct localization of Wnt7b, Notum, and Wnt5a in developing trachea supports a role for Notum in regulating the levels and the types of activity triggered by Wnt7b and Wnt5a. As previously shown, Notum can attenuate the activation of Top Flash by the canonical ligand Wnt3a (Gerhardt et al., 2018) Kakowaga et al, 2016). Similarly, Notum reduces Wnt5a-induced activation of ATF2 (Ohkawara and Niehrs, 2011; Wallkamm et al., 2014) (Fig 1c) and AP1 promoters (data not shown) (Nishita et al., 2010). These two transcription factors are part of the Wnt non-canonical/Planar Cell Polarity pathway and are downstream of JNK (Fig 1c). These results suggest that Notum can deactivate ligands from canonical and non-canonical Wnt signaling.

**Figure 1:**
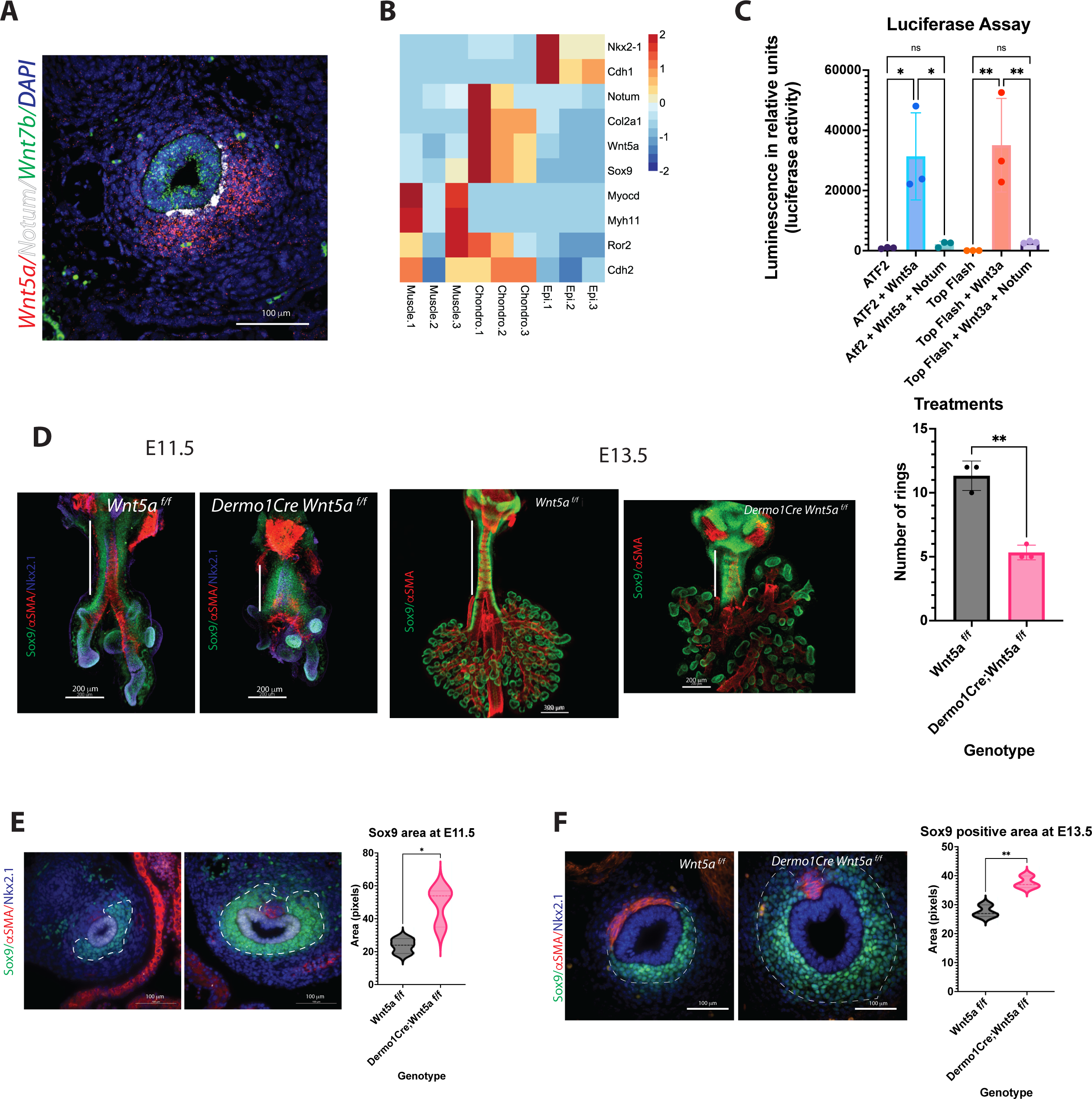
A) RNA in situ hybridization of E13.5 cross section depicting the localization pattern of Wnt non-canonical ligand *Wnt5a* (red) in the ventrolateral region of the trachea mesenchyme, and canonical ligand *Wnt7b* (green) expressed in the epithelium of the trachea. *Notum* (white) is located in the mesenchymal, subepithelial region in between *Wnt5a* and *Wnt7b* areas of expression. DAPI (blue) was used to stain the nuclei. 40x image. B) Heatmap of RNAseq from cell sorted tracheal cells isolated at E13.5. Epi. antibody stained EpCAM (+) cells, Muscle. endogenous SMA-eGFP (+) cells, Chondro. EpCAM (-), SMA-eGFP (-) double negative cells. *Notum* and *Wnt5a* are expressed in chondroblast cells, identified by the expression of chondrogenic genes: *Col2a1* and *Sox9*. Otherwise absent in muscle and epithelial cells. The expression of typical markers in the three cell lineages, confirms a good result of the cell sorting strategy. C) Luciferase assay performed on NIH3T3 cells transfected with ATF2 or TOP Flash reporter constructs. Co-transfection of Wnt5a expression plasmid increases the luciferase activity by the ATF2 construct. In case of Wnt3a expression plasmid, it increases the luciferase activity by the TOP Flash construct. In both cases, when Notum expression plasmid is also added, this luciferase induction is abrogated. In this in vitro assay, Notum can dampen the activities of canonical (Wnt3a) and non-canonical (Wnt5a) ligands. n=3 for each treatment group. * p<0.05, ** p<0.01, ns non-significant statistical differences. One-way ANOVA D) <u>Left and center panel:</u> Wholemount immunofluorescence (IF) of dissections of larynx-trachea-lungs at E11.5 and E13.5 of control (Wnt5a ^f/f^) and Wnt5a mutants (Dermo1cre; Wnt5a ^f/f^). Sox9 (green) is expressed in chondroblasts and epithelial cells of the distal lung. αSMA (red) is localized in muscle cells of the dorsal trachea and bronchi. This expression pattern is maintained in Wnt5a mutants compared to controls, but the trachea is shorter in length (white vertical full line) with defects in lung lobation noticeable by E13.5. At E11.5, there is no evidence of ring formation as noticed by the Sox9 (+) continuum observed in the trachea mesenchyme. At E13.5, Sox9 (+) cells in the trachea appear more defined in rings for Wnt5a mutants than controls. Dorsal trachea muscle cells (αSMA (+) show defects in orientation, particularly towards the proximal region, closer to the larynx. Scale bars: 200 um for E11.5 samples, 300um for the E13.5 control image, and 200um for the E13.5 Wnt5a mutant image. <u>Right panel:</u> Reduction in the tracheal ring number by E14.5 in mutans vs controls. n=3 for each genotype. E) IF of cross sections of control and Wnt5a mutant at E11.5. Chondroblasts are Sox9 (+) (green), muscle cells are αSMA (+) (red), and respiratory epithelial cells are Nkx2.1 (+) (purple). In wnt5a mutants Sox9 (+) area is thicker in the mutant than the control. N=3 p<0.05 F) IF of cross sections of E13.5 embryos. Sox9 (+) (green) area (delimited by a dashed white line) is thicker in Wnt5a mutants compared to controls, as quantified by Sox9 positive staining. Sox9 expression also extends dorsally. Muscle cells (αSMA (+), red) are organized differently in Wnt5a mutants vs controls. DAPI (blue) was used to stain the nuclei. 40x images. N=3 p<0.01 Cross-sectional slices of the trachea were consistently obtained at the location of the heart.

To better understand Wnt5a’s putative role in cartilaginous ring formation, we deleted *Wnt5a* from the splanchnic mesoderm using *Dermo1Cre*. Analysis of whole mount images and cross sections of *Dermo1Cre;Wnt5a^f/f^* trachea demonstrated the deficiencies in tracheal elongation as early as E11.5, and reduced number of cartilaginous rings by E14.5, in agreement with published studies (Kishimoto et al., 2018; Li et al., 2005). Strikingly, we detected that the cartilaginous rings appear more defined (Fig 1d), with a larger field of Sox9 positive cells (chondroblasts) in the Wnt5a deficient trachea as early as E11.5 and had abnormal organization of the trachealis muscle noticeable by E13.5 (Fig 1d,f).

Taken together, Notum can attenuate the non-canonical induced Wnt5a activity in vitro. In vivo, Wnt5a is required for proper tracheal elongation, cartilaginous ring number, and shape of the tracheal cartilage.

### Precocious mesenchymal condensation in Wnt5a is associated with JNK and Ca2++ signaling

Our previous studies determined that Notum delays the mesenchymal condensation, resulting in thinner and mis-patterned cartilage (Gerhardt et al., 2018). Since our observations at E13.5 suggest a precocious condensation in the Wnt5a mutant trachea, we performed a serial analysis of the mesenchymal condensation process from E11.5 to E14.5 in Wnt5a and Notum deficient tracheas. At E11.5, when tracheoesophageal separation is completed, Sox9+ cells are distributed in a continuous stripe for all genotypes analyzed. Interestingly, the Wnt5a deficient trachea is already shorter than the control and the Sox9+ area is thicker than the control (Fig.1f). Similarly, at E12.5, no presence of aggregation of Sox9+ cells are detected, indicating that no mesenchymal condensations are visible; however, as early as E13.0, we noticed the formation of mesenchymal nodules in Wnt5a deficient trachea, which become clearly defined by E13.5 (Fig 2a, b). Contrasting this finding and concordant with our published studies, mesenchymal condensations are delayed until E14.5 in the Notum deficient tracheas (Fig2 a,b,c). To further understand the underlying mechanism by which Wnt5a and Notum could affect the timing of the mesenchymal condensation, we performed an air-liquid interface (ALI) culture using trachea-lung tissue isolated from E12.5 *Sox9KIeGFP* mice. Tissue treated with ABC99, a Notum inhibitor, shows no sign of cartilaginous mesenchymal condensations at 42 hours compared to control (Fig.3a and Supplement, Video 2, compared to control Video 1). The treatment recapitulates the in vivo findings observed in the Notum deficient trachea (Fig 2a). Wnt5a has been shown to act via Ror1/2 receptors activating β-catenin independent Wnt signaling pathways (Kishimoto et al., 2018), including planar cell polarity (PCP) and Calcium dependent signaling (Chae and Bothwell, 2018; Kikuchi et al., 2011). Thus, we pharmacologically inhibited both pathways in E12.5 *Sox9KIeGFP* trachea-lung tissue. Inhibition of non-canonical Calcium dependent Wnt signaling with KN93 (calmodulin kinase (CamK) inhibitor) or inhibition of the PCP pathway with JNK inhibitor cause earlier condensation at 24 hours of incubation compared to controls (Fig 3a,b) (Supplement, Videos 3 and 4). These findings recapitulate the in vivo observations in the Wnt5a deficient trachea and indicate that the Ca2++ and the JNK non-canonical Wnt pathways are required for regulation of timely mesenchymal condensation and a possible mechanism downstream of Wnt5a.

**Figure 2:**
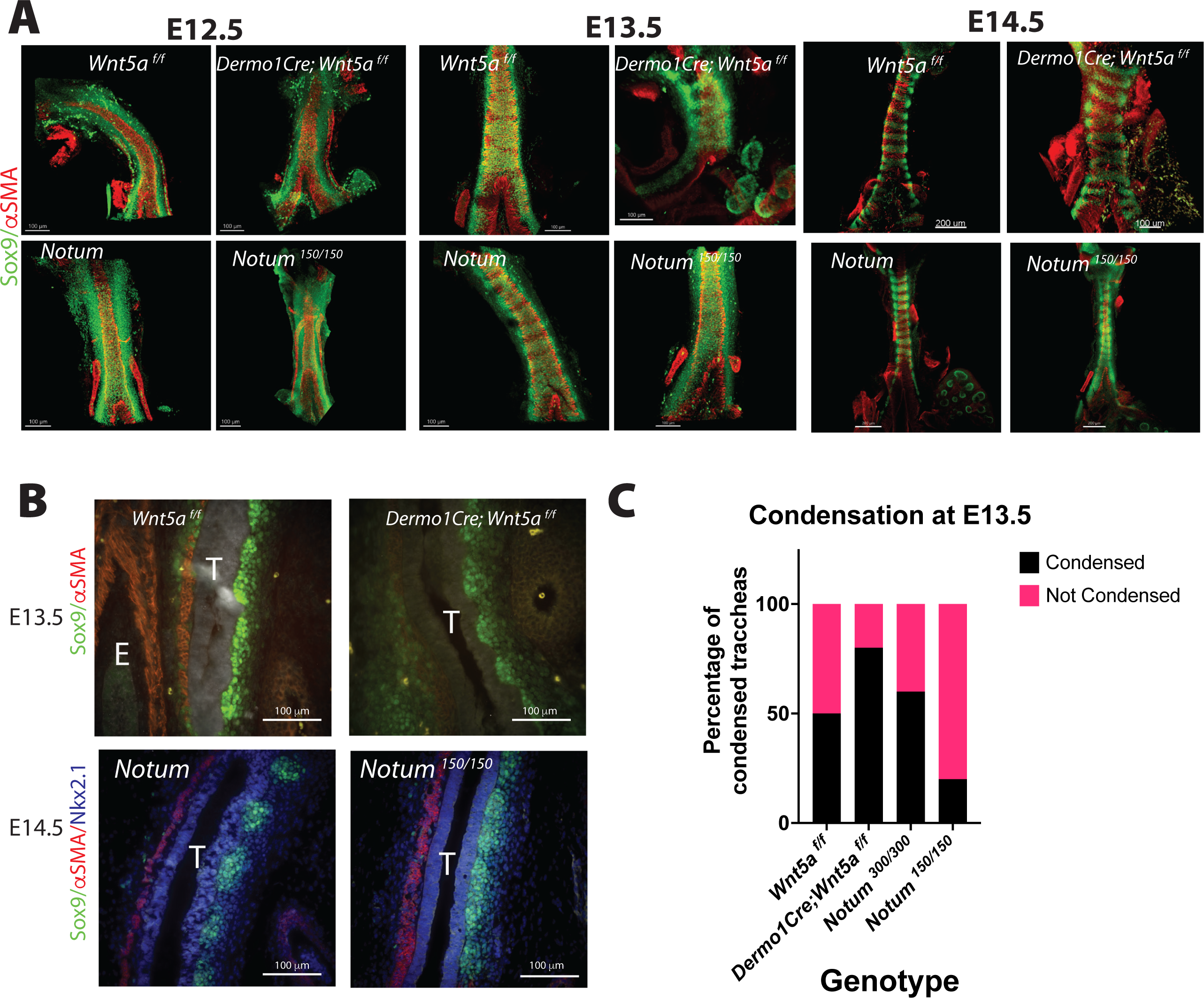
A) Wholemount IF of E12.5 to E14.5 dissected tracheas. In this tissue Sox9 is expressed by ventrolateral chondroblasts (green) and αSMA by the dorsal trachealis muscle cells and surrounding blood vessels (red). By E12.5 controls (Notum and Wnt5a^f/f^) show a continuum of Sox9 expression that starts to condensate at E13.5, showing a defined separation in rings by E14.5. Top panel: Wnt5a mutant tracheas (Dermo1cre; Wnt5a ^f/f^) although shorter in length show a similar phenotype at E12.5 than controls, but at E13.5 rings are already visible with a higher density of Sox9 (+) cells. By E14.5 rings in the Wnt5a mutant are completely formed with morphological alterations compared to controls. <u>Bottom panel:</u> Notum mutants (Notum ^150/150^) also present a Sox9 homogeneous expression at E12.5. Differently to their control counterparts and Wnt5a mutants, by E13.5 there is no sign of mesenchymal cell condensations. Even by E14.5 Notum mutants show a lagging and altered cartilage ring formation and a stenotic trachea compared to controls. Scale bars: 100um for E12.5 and E13.5 images, 200um for E14.5 images (except Wnt5a mutant, 100um). B) IF in longitudinal sections of E13.5 embryos. Sox9 (+) (green) cell condensations are more defined in Wnt5a mutants than controls. αSMA (red) is visualized in the dorsal aspect of the trachea (T), and in the esophagus (E). In contrast, at E14.5 condensations are not formed in Notum deficient tracheas as depicted by the Sox9 staining. C) In a frequency analysis from the wholemount IF stainings we observe that by E13.5, the percentage of condensed tracheas (black bars) is higher in Wnt5a mutants than controls. The opposite behavior is present in Notum mutants, which have a lower percentage of condensed tracheas than Notum controls. n= 5-13, for each genotype.

**Figure 3:**
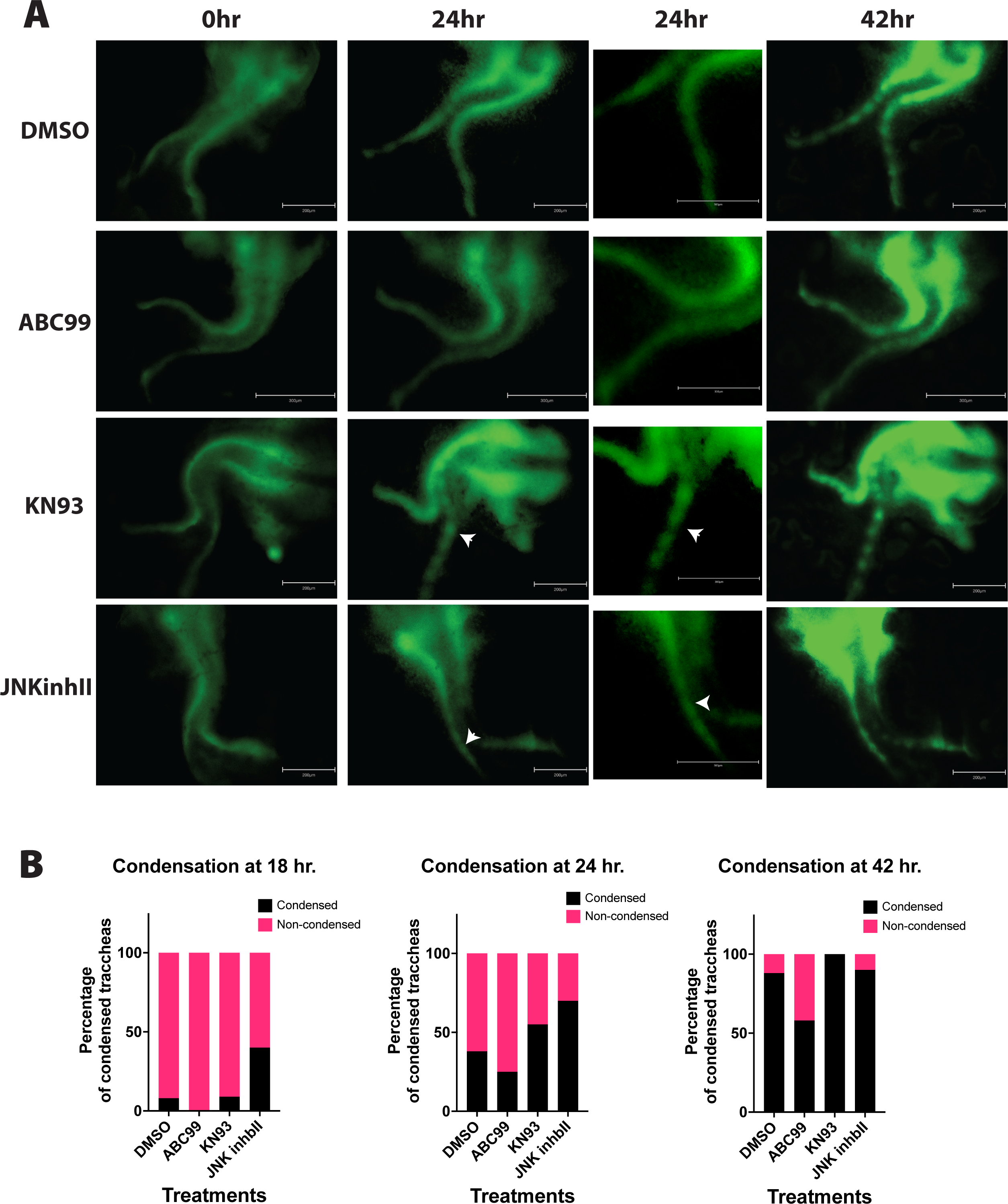
A) Timelapse imaging of larynx-trachea-lung dissected tissue from E12.5 embryos grown ex vivo in Air Liquid Interphase (ALI). Endogenous expression of Sox9KIeGFP was used to visualize the moment of cell condensation that precedes the formation of cartilage. Condensations in ex- vivo explants were defined as SOX9 GFP+ cells that formed aggregates and became segmented as time progressed. Vehicle treated tissue (DMSO) shows signs of condensation at 42 hours of incubation. This phenomenon is absent when Notum is chemically inhibited (ABC99). Inhibitors of the Wnt non-canonical pathway, Calmodulin Kinase inhibitor (KN93) and JNK inhibitor (JNKinhII), induce earlier signs of Sox9KIeGFP (+) condensations at 24 hours (white arrows), earlier than vehicle treated samples. Scale bar 300 um. Digital zoom for images at 24hours is shown, scale bar 150 um. B) In a frequency analysis from the ALI cultured samples, we observed a trend towards a higher percentage of condensations as early as 18 hours in JNKinhII treated samples compared to vehicle treated samples. At 24 hours KN93 and JNKinhII show higher percentages of tracheal condensations than vehicle treated samples. ABC99 treated samples show a reduced percentage of tracheal condensations at 18, 24 and 42 hours compared to vehicle or other treatments. n= 7- 16 for each treatment.

### Cell mechanisms mediating cartilaginous condensation are altered in Wnt5a deficient and Notum deficient tracheas

The mesenchymal condensation process requires precise cellular behaviors leading to the compaction of mesenchymal cells (Klumpers et al., 2014). Thus, we sought to investigate whether essential mechanisms mediating mesenchymal cell condensation are affected in Wnt5a and Notum mutants. Evaluation of the migratory ability of E13.5 Wnt5a deficient mesenchymal cells in a wound healing assay revealed Wnt5a knockout cells moved slightly slower compared to the control. At 24 hours, the Wnt5a knockout cells average a 40% closure of the wound area compared to almost 50% for the control; however, no statistical differences are detected.

On the other hand, evaluation of the wound-healing assays showed a trend for the Notum knockout cells moving faster than controls (Fig. 4a). Since migration of condensing cells is a local process of directional movement induced by cues present in the mesenchymal ECM, we investigated whether the directional migration toward fibronectin is affected in E13.5 cells of Wnt5a, and Notum deficient tracheal mesenchymal cells in comparison to their respective controls. Notably, the Notum deficient cells had the highest directional migratory ability in this haptotaxis assay, while the Wnt5a mutant cells did not differ from the control (Fig. 4b). Compaction of mesenchymal cells requires changes in cell-cell and cell-substrate adhesion; we tested the adhesion of E13.5 tracheal primary mesenchymal cells as micromasses. When seeded as micro masses, Notum knockout cells show a trend towards reducing cell adhesion compared to controls. Wnt5a knockout cells show a slight increase in cell adhesion compared to controls; however, no significant differences are observed among the genotypes (Fig 4c). Since mesenchymal condensations occur prematurely in Wnt5a deficient tracheas (Fig 2 and 3), with cells already condensed by E13.5, we analyzed whether cell-cell adhesion molecules are prematurely expressed in Wnt5a mutant tracheas. Thus, we performed N-cadherin immunofluorescence stainings at E11.5 (Fig 4 d) and E13.5 (not shown). N-cadherin is an essential molecule mediating the adhesion of condensed cells. We observed an increased expression of N-cadherin in *Dermo1Cre;Wnt5a^f/f^*, at E11.5, validated by the quantification of the staining in regions overlapping with Sox9 staining. Further, Sox9 staining depicts Sox9+ cells appearing more compacted as opposed to the loose appearance of the Sox9+ cells in the *Wnt5a^f/f^* trachea (Fig 4d). Taken together, changes in cell behavior support the differences in the timing of chondrogenesis in the Wnt5a knockout and Notum knockout tracheas. The increased movement of Notum knockout cells at E13.5 likely explains the delayed chondrogenesis at E14.5. In contrast, the slower cell movement of Wnt5a knockout cells and the precocious expression of N-cadherin support the earlier chondrogenesis, in mutant Wnt5a tracheas.

**Figure 4:**
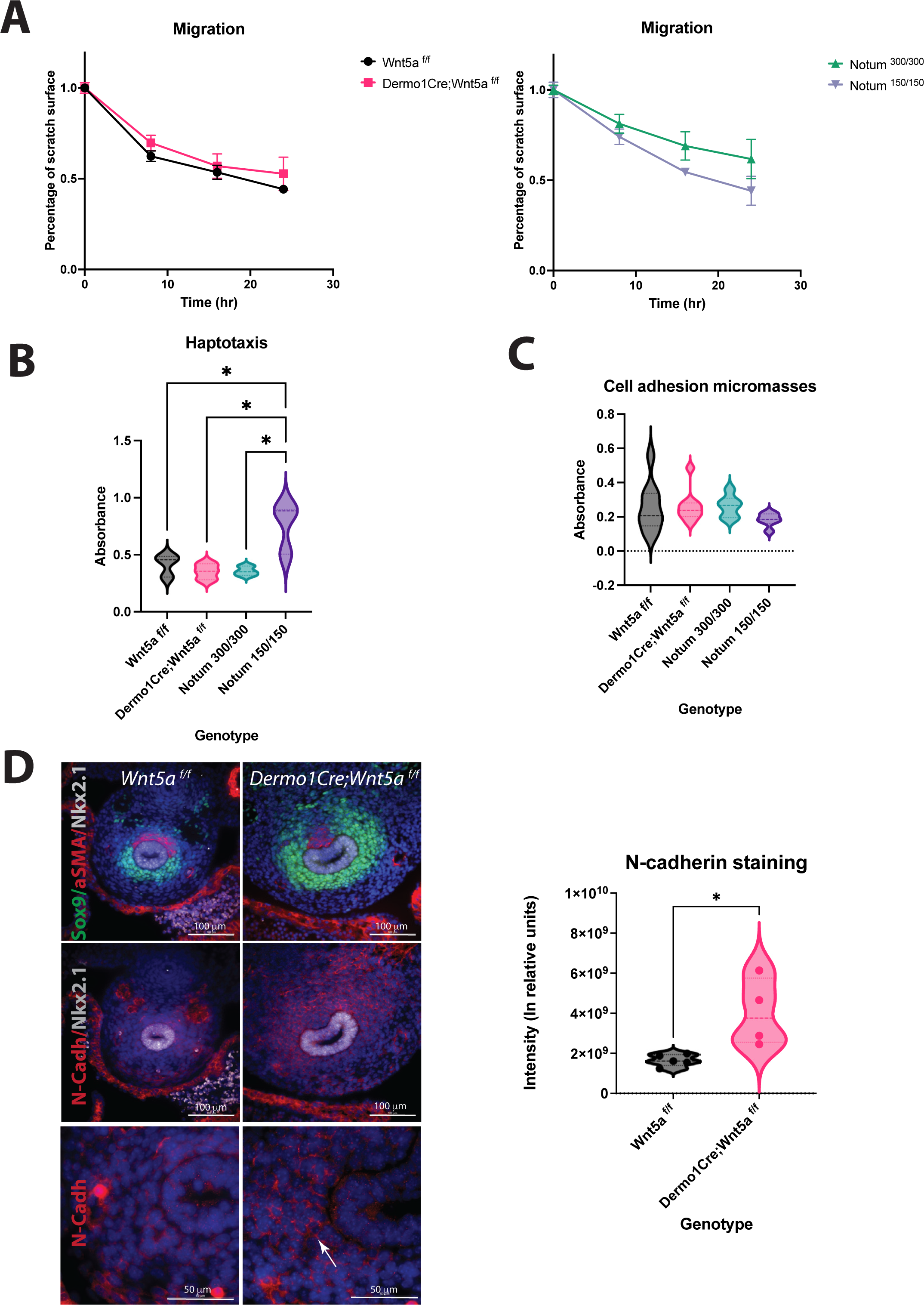
A) Left <u>panel:</u> Migration assays of E13.5 primary mesenchymal tracheal cells show no statistical differences in cell motility between Wnt5a controls (*Wnt5a ^f/^*^f^, black circles) and Wnt5a mutants (*Dermo1cre; Wnt5a ^f/f^*, pink squares). <u>Right panel:</u> There is a trend towards a higher motility of Notum mutants (Notum ^150/150^, purple triangles) compared to Notum controls (Notum ^300/300^, green triangles) at 16 and 24 hours. n=3 for each genotype. B) Haptotaxis assays of E13.5 primary mesenchymal tracheal cells show an increase in directional cell migration towards fibronectin for Notum mutants (purple) compared to Notum controls (green) and other genotypes. Wnt5a mutants (pink) do not show a difference in directional cell migration compared to Wnt5a controls (black). n=3 per genotype. * p<0.05 One- way ANOVA. C) Cell adhesion in micromasses from E13.5 primary mesenchymal tracheal cells showed no differences in the genotypes analyzed. n=6-9 per genotype. D) Immunofluorescence stainings of E11.5 cross sections. <u>Left panel, top two images:</u> Sox9 (+) (green) area is thicker in Wnt5a mutants compared to controls. Sox9 expression also extends dorsally. Muscle cells (αSMA (+), red) are organized differently in Wnt5a mutants vs controls. Respiratory epithelium identity is maintained (Nkx2.1, white). 40x images. <u>Left panel, two images</u> <u>in the middle:</u> N-cadherin (N-cad, red) is increased in Wnt5a mutants compared to controls. Respiratory epithelium is stained with anti-Nkx2.1 (white). 40x images. <u>Left panel, bottom two</u> <u>images:</u> N-cadherin expression is stronger in the cell-cell interphase of dorso-lateral mesenchymal cells. 100x oil immersion images. DAPI (blue) was used to stain the nuclei. <u>Right panel:</u> N-cadherin intensity quantification of the experiments presented in (E) shows an increase for Wnt5a mutans (pink) compared to controls (black). n=4-5 per genotype, p=0.01 T test.

### Molecular signatures in Wnt5a deficient trachea indicate premature activation of a gene expression network leading to a chondrogenic process

Based on changes in cell-cell adhesion and the premature condensation observed in the Wnt5a deficient trachea, we tested the hypothesis that Wnt5a prevents premature mesenchymal condensation via modulation of chondrogenic gene expression. The transition from E11.5 to E13.5 is critical for tracheal development. During these stages a distinct tracheal tube is formed consisting of Nkx2.1+ epithelium surrounded by its unique mesenchyme. By E13.5 trachealis muscle will be organized in the dorsal side, while the mesenchymal condensations that give rise to tracheal cartilage are initiated in the ventral mesenchyme. To define molecular mechanisms explaining the cellular behaviors observed in the Wnt5a deficient trachea, we first performed RNA seq analysis of E11.5 vs. E13.5 wild type tracheal tissue to determine changes in gene expression occurring between these critical developmental stages (Fig 5a). Genes differentially regulated and augmented at E13.5 include those encoding ECM molecules *Acan* and *Col9a1*, the transcription factor *Barx1*, and signaling molecules such as *Gdf5, which is* essential for cartilage formation (Fig 5a). Among biological processes enriched at E13.5, those related to skeletal development, ECM organization, and cartilage development were more significant. Top induced pathways include “ECM organization” and signaling pathways, including FGF and TGF-beta signaling pathways (Fig 5b). Next, we performed whole bulk RNA seq on E11.5 tracheas of *Wnt5a^f/f^* and *Dermo1Cre; Wnt5a^f/f^* mice. We selected this stage because the trachea is fully separated from the esophagus, and it precedes the morphological changes associated with mesenchymal chondrogenesis. Furthermore, we inferred from our previous findings that Wnt5a deficient tracheas may start the chondrogenic process precociously. An unbiased gene expression profiling analysis of Wnt5a mutant vs control shows a premature upregulation of chondrogenic genes by E11.5, including *Col2a1*, a marker of chondrogenesis. We also detected that genes that are normally upregulated at E13.5 in control trachea, i.e., *Gdf5, Barx1, Col9a1*, are also prematurely induced in the Wnt5a mutant, at E11.5, supporting the notion that the chondrogenic program is precocious in the absence of Wnt5a (Fig 5c). In agreement with this finding, biological processes enriched in the mutant Wnt5a trachea include those such as cartilage development, chondrocyte differentiation, and regulation of the Wnt signaling pathway (Fig.5d). Since cell compaction induces specific cell differentiation processes in mesenchymal condensations, we examined expression levels of genes encoding transcription factors with a cis-regulatory region binding capable of regulating chondrogenesis. As anticipated, transcription factors such as *Barx1, Osr2, Hoxc5,* and *Tbx18* are differentially regulated in E13.5 wild type tracheas. Remarkably these genes are also upregulated at E11.5 in the *Dermo1Cre; Wnt5a^f/f^* trachea (Fig 5e). The data support a model whereby the expression of Wnt5a modulates the timely activation of the chondrogenic process in tracheal mesenchyme by controlling gene expression associated with chondrocyte differentiation.

**Figure 5:**
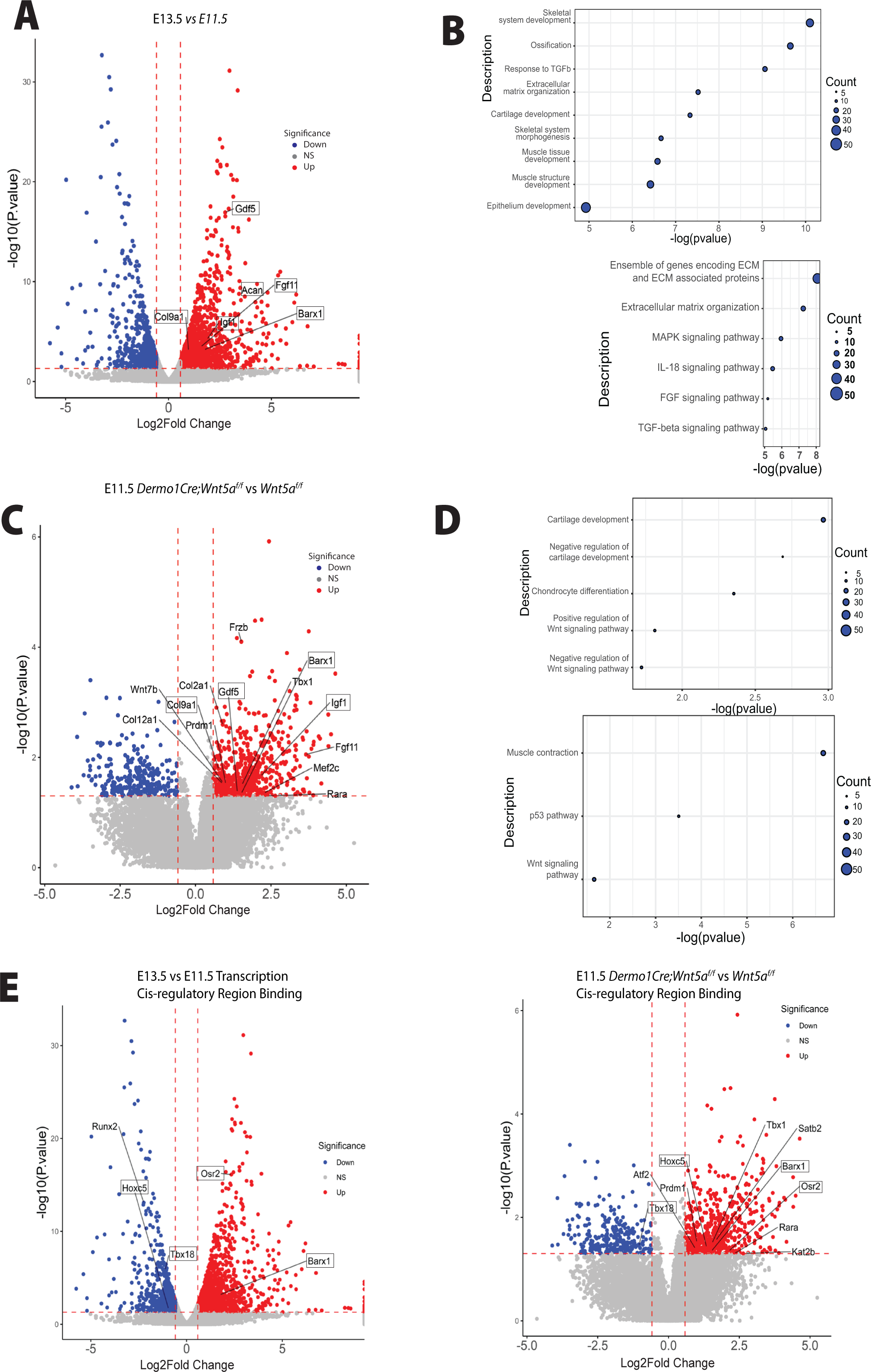
A) Volcano plot depicting E13.5 vs E11.5 genes of interest which have been differentially regulated. Specifically, *Gdf5, Col9a1, Igf1, Barx1*, and *Acan* are upregulated at E13.5. Blue: downregulated, red: upregulated genes. B) Bubble plots depicting E13.5 vs E11.5 biological processes and pathway enrichments with significance. C) Volcano plot depicting E11.5 *Dermo1Cre;Wnt5a^f/f^* vs *Wnt5a^f/f^* genes of interest which have been differentially regulated. Specifically, Gdf5, Col9a1, Igf1, and Barx1 are upregulated at E11.5 after the conditional deletion of Wnt5a. Blue: downregulated, red: upregulated genes. D) Bubble plots depicting E11.5 *Dermo1Cre;Wnt5a^f/f^* vs *Wnt5a^f/f^* biological processes and pathway enrichment with significance. E) Volcano plot depicting differentially regulated genes of interest with cis-regulatory region binding on chondrogenic genes of E13.5 vs E11.5 (left) and E11.5 *Dermo1Cre;Wnt5a^f/f^* vs *Wnt5a^f/f^* (right). *Osr2* and *Barx1* are upregulated while *Tbx18* and *Hoxc5* are downregulated in the E13.5 vs E11.5 dataset. *Tbx18*, *Hoxc5*, *Osr2*, and *Barx1* are upregulated in the E11.5 *Dermo1Cre;Wnt5a^f/f^* vs *Wnt5a^f/f^* dataset. Blue: downregulated, red: upregulated genes.

### Coordinated regulation of the chondrogenic program by Wnt ligands

Our in vitro studies demonstrated that Notum modulates Wnt5a-induced activity (Fig 1c). On the other hand, deletion of Wnt5a accelerates the chondrogenesis program, a phenotype somewhat opposed to the Notum deficient trachea, wherein chondrogenesis is delayed (Fig 2). We reasoned those molecular signatures contributing to the premature condensation observed in the *Dermo1Cre;Wnt5a ^f/f^* trachea might be downregulated in the Notum model wherein condensation is delayed. To test this concept, we performed gene expression analysis of chondroblast cells of *Notum* deficient tracheas at E13.5, a developmental stage when cartilaginous mesenchymal condensation is already initiated in the wild-type trachea (Fig 6a). Supporting our published studies, the dataset reveals increased expression of Wnt signaling related genes such as *Axin2, Ctnnb1, Lef1* (Fig 6a). Our RNA seq data identified several genes associated with Wnt/b-catenin independent signaling pathway differentially regulated in Notum deficient tissue, including Dvl1 Dvl1 is an intracellular transducer mediating different non-canonical Wnt pathways (Sharma et al., 2018). Variants in DVL, together with variants in WNT5a and ROR2, have been associated with Robinow syndrome, a condition associated with skeletal malforamtions (Konopelski Snavely et al., 2023). We also detected upregulation of Fat1 a gene associated with PCP, encoding an atypical cadherin playing a critical role in directed cell migration and cell-cell contact (Peng et al., 2021) (Fig 6a). Despite these changes in gene expression associated with non-canonical Wnt signaling supporting a potential increase in the pathway activity, we did not observe a significant increased expression of *Wnt5a* in *Notum* deficient tracheas (Supplement Fig2).

In *Notum* mutant samples, we did observe differential expression of genes mediating mesenchymal cell differentiation, such as *Fgf10, Bmp3, Gata4, Col2a1*, and an increased expression of genes influencing cartilage formation including *Sox5, Msx2*, and *Arid5a (Fig.6b)*. Further, genes involved in cartilage development, condensation, and mesenchymal cell differentiation were largely induced based on the functional enrichment analysis. Corroborating ours and others published findings, pathways enriched included Wnt signaling, Notch, and TGFbeta (Fig 6c). We then compared the differentially regulated genes in response to Wnt5a deletion vs. Notum deletion and detected a discrete list of commonly regulated genes after deleting either Wnt5a or Notum. For some of these genes, the RNA seq predicted opposite regulation; however, other genes, such as Col2a1, were upregulated in both datasets (Fig 6d). A delayed chondrogenic program in the Notum model could partially explain the data. To validate the findings of the RNAseq studies, we performed RNA in situ hybridization in samples of E11.5 *Wnt5a ^f/f^* (control) and *Dermo1Cre; Wnt5a ^f/f^* (mutant) and E13.5 *Notum ^300/300^* (control) and *Notum ^150/150^* (mutant) (Fig.6e). We first tested expression of Gdf5, which encodes a secreted growth factor promoting cartilage and bone development. As predicted by the analysis of developmental differential gene expression (Fig.5a,c), levels of expression of *Gdf5* were low at E11.5 as demonstrated by the small number of transcripts detected in Wnt5a controls when compared to E13.5 Notum control trachea. On the contrary levels of *Gdf5* transcripts were highly increased in the *Wnt5a* deficient trachea, resembling the E13.5 expression pattern (Fig 5.ad). No differences in expression levels of *Gdf5* are detected in *Notum* control and *Notum* deficient trachea. *Igf1* transcripts, encoding a growth factor necessary for postnatal lung alveologenesis (Gao et al., 2022), were increased in the dorsal and to some degree the ventral mesenchyme of the *Wnt5a* deficient trachea and with a similar expression pattern seen at E13.5 control trachea. No significant differences in *Igf1* expression were observed between control and *Notum* deficient trachea. Concordant with the RNA seq data, *Gata4* transcripts, encoding a transcription factor required for muscle development and pulmonary lobar morphology (Ackerman et al., 2007), are decreased in *Wnt5a* deficient trachea, and increased in *Notum* deficient trachea. *Osr2* transcripts (Rankin et al., 2012) are observed in an opposite pattern, wherein transcripts are increased in the epithelium of the Wnt5a deficient trachea, while decreased in the epithelium and ventral distal mesenchyme of the *Notum* deficient trachea.

Since the RNA seq data suggest that the transcriptional program controlled by Wnt5a and Notum are likely independent and effects on gene expression are likely indirect, we tested whether increased levels of Wnt5a could account for delayed chondrogenesis as observed in Notum deficient trachea. For this purpose, we performed ALI culture of explants isolated at E12.5 and incubated with Wnt5a conditioned media or in presence of the Notum inhibitor ABC99. Trachea lung explants were harvested at 42 hrs, time when condensations were observed in control tracheal lungs (Fig 3.a). After collecting the tissue, explants were processed for whole mount immunofluorescence to determine the expression patterns of Sox9 and αSMA. As anticipated, in DMSO treated tracheas Sox9 positive cells condensed forming clear bands of cell aggregates. Strikingly, tracheas incubated in presence of Wnt5a conditioned media do not condense or condensation was lagging when compared to control DMSO treatment. The finding recapitulates the observed lack of mesenchymal condensation seen after treatment of the trachea- lung explants with ABC99 (Fig.7a). The fact that treatment with conditioned Wnt5a media caused the delay of mesenchymal condensations made us consider whether in the Notum mutant the delayed mesenchymal condensation is caused by increased levels of Wnt5a. We generated embryos of genotype *Dermo1Cre; Wnt5a ^f/wt;^ Notum ^150/150^* and evaluated the presence of mesenchymal condensations at E13.75 in whole mount immunofluorescence (Fig.7b). The Sox9 staining revealed that condensations were visible in the control embryos (*Wnt5a^f/wt^;Notum ^300/150^*, *Dermo1Cre; Wnt5a^f/wt^; Notum^300/150^*, or *Dermo1Cre; Notum^300/150^*) and were not detected in the Notum deficient embryos (*Wnt5a ^f/wt;^ Notum^150/150^*). Strikingly, in embryos of genotype *Dermo1Cre; Wnt5a ^f/wt;^ Notum ^150/150^*, condensations were visible as determined by the Sox9 staining. Taken together, the data support a model wherein Wnt5a plays a critical role in controlling the timing of the cartilaginous mesenchymal condensation while Notum attenuates the levels of Wnt5a activity.

**Figure 6:**
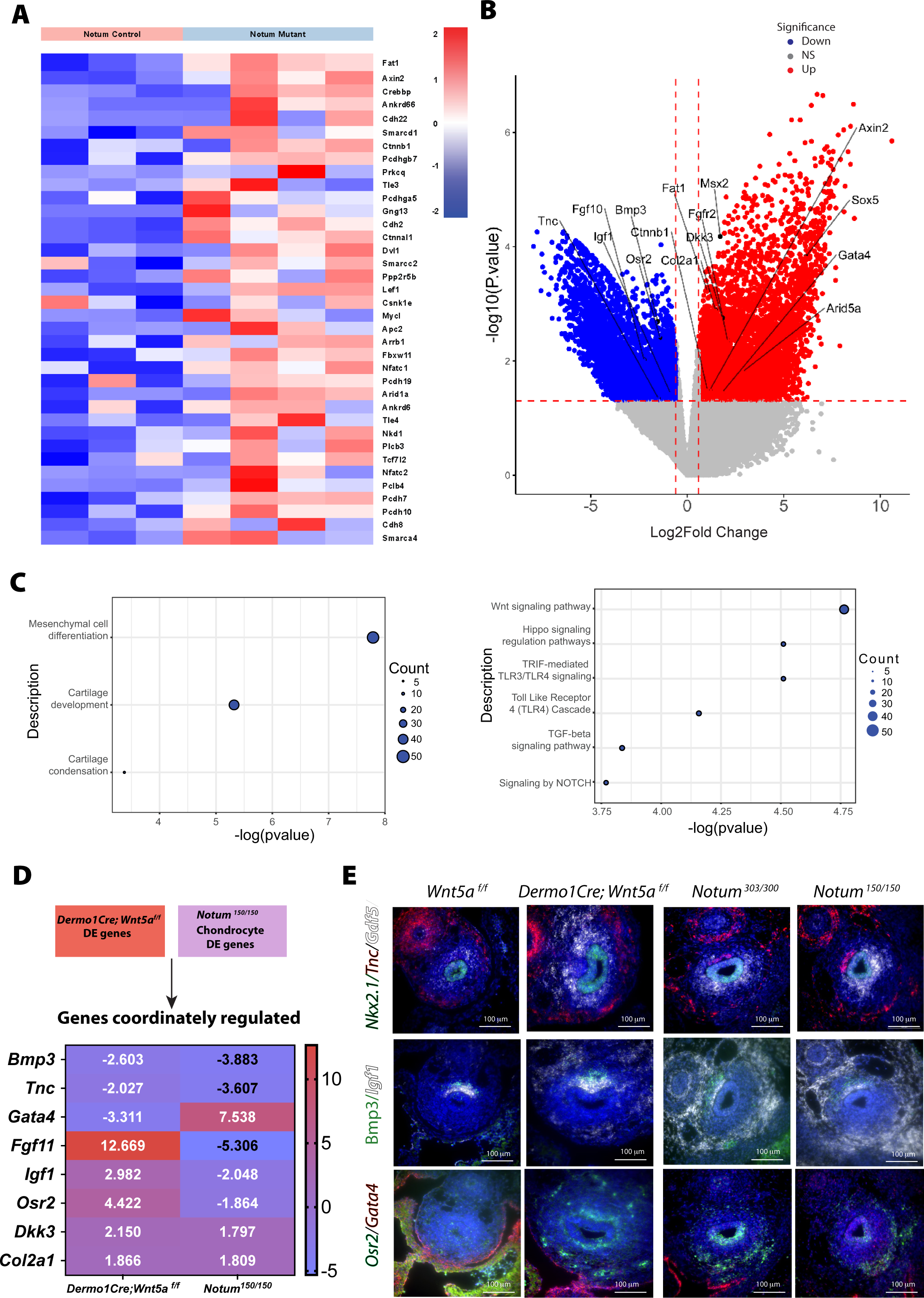
A) Heat map demonstrates differential expression of Wnt signaling pathway genes among Notum control (Notum ^300/300^) and mutant (Notum ^150/150^) cells of the tracheal chondroblasts at E13.5. The dataset reveals increased expression of *Axin2, Ctnnb, Dkk3, and Dvl1*. B) Volcano plot depicting Notum mutant (*Notum^150/150^)* vs control (*Notum^300/300^*) genes of interest which have been differentially regulated in tracheal chondroblasts. Blue: downregulated, red: upregulated genes. C) Bubble plots depicting Notum biological processes and pathway enrichments with significance are shown. D) Heatmap of common genes differentially regulated in cells of *Dermo1Cre;Wnt5a^f/f^* and *Notum^150/150^* chondrocytes. *Gata4* appears more upregulated in Notum cells while *Fgf11* and *Osr2* appear more upregulated in *Dermo1Cre;Wnt5a^f/f^* cells. Blue: downregulated, red: upregulated genes. E) RNA in situ hybridization depicts localization of transcripts for Nkx2.1, *Tnc, Gdf5, Bmp3, Igf1*, *Osr2*, and *Gata4* in transverse sections of E11.5 *Wnt5a^f/f^*, *Dermo1Cre;Wnt5a^f/f^*, E13.5 *Notum^303/300^*, and *Notum^150/150^* tracheas. *Osr2* appears to be more abundant in Wnt5a mutant tracheas and *Gdf5* appears more abundant in E11.5 Wnt5a mutant tracheas and in E13.5 vs E11.5 controls which validates previous RNAseq findings.

**Figure 7:**
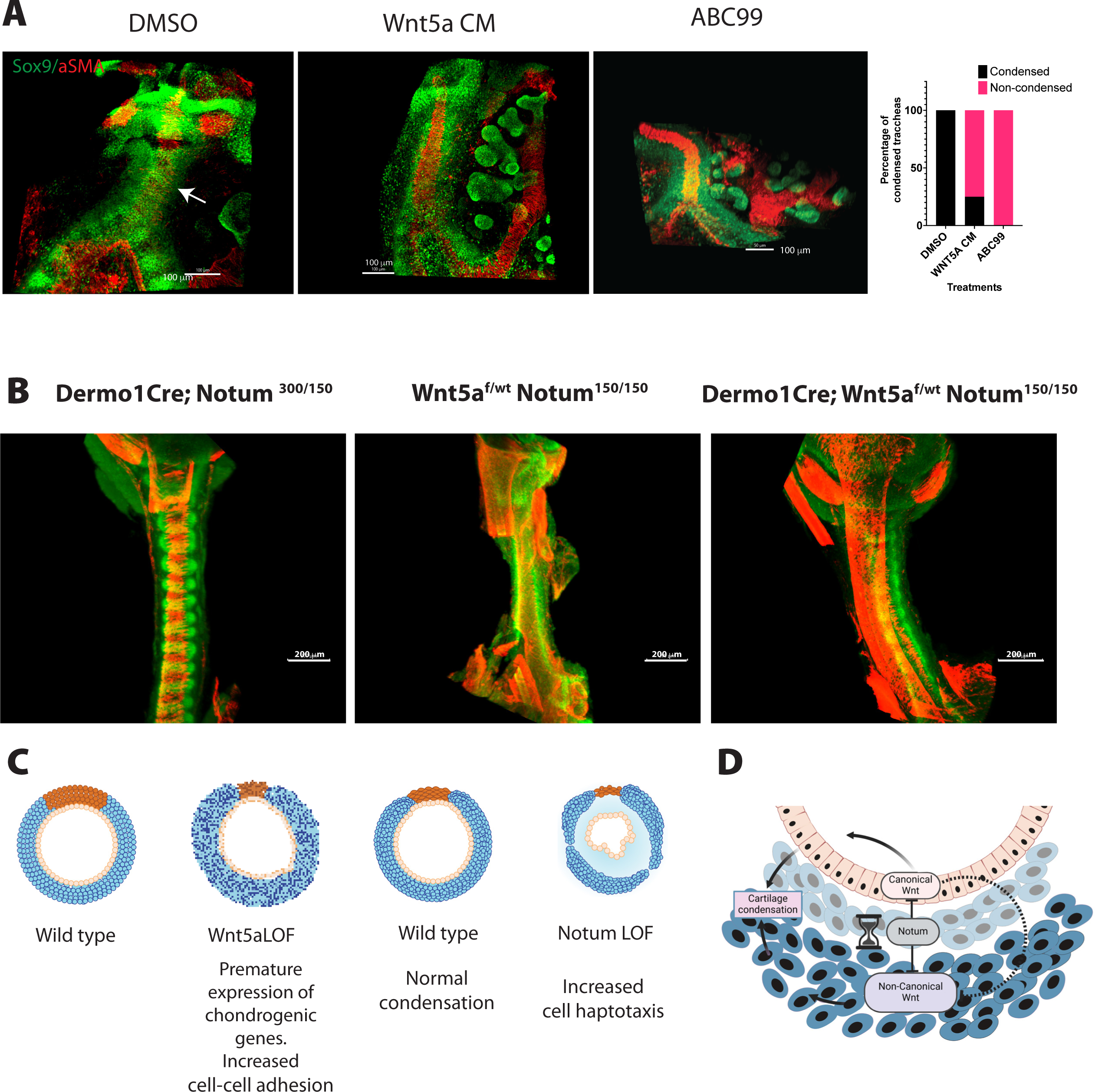
A) Whole mount immunofluorescence imaging of wild type tracheal lung explants are depicted after 42 hrs after ALI cultures. Sox9 (green) is expressed in chondroblasts and epithelial cells of the distal lung. αSMA (red) is localized in muscle cells of the dorsal trachea and bronchi. In vehicle treated tracheas, condensations were clearly observed at 42 hrs (arrow DMSO). In contrast, addition of Wnt5a conditioned media in media culture prevented the formation of cartilaginous mesenchymal condensations, recapitulating the phenotype obtained after treatment with ABC99, a Notum inhibitor. Frequency analysis depicting the effects of the treatments is shown to the right of the whole mount images. N=4-8 per treatment. B) Whole mount immunofluorescence of E13.75 embryos of a Notum Wnt5a allelic series demonstrating the rescue of the timing of mesenchymal condensations by decreasing the Wnt5a expression in Notum deficient background (*Dermo1Cre; Wnt5a^f/wt^; Notum ^150/150^*) as demonstrating by the Sox9 staining (green). Images are representative of embryos from three different litters. C) The cartoon summarizes the effects of deletions of Wnt5a and Notum in mesenchymal cell condensation and cartilage. Deletion of Wnt5a (Wnt5a LOF) causes premature condensation of chondroblasts associated with increased cell-cell addition and the premature expression of genes promoting mesenchymal condensations at E11.5. On the other hand, after deletion of Notum (Notum LOF), mesenchymal condensations are delayed in association with increased cell directional migration at E13.5. In both models, the normal morphology of the cartilage is disrupted. D) Proposed model integrating Notum, Non-canonical and Canonical Wnt signaling. Notum balances Wnt signaling in developing trachea by directly inhibiting the activity of both Wnt5a- mediated non-canonical and canonical Wnt signaling. Indirectly, by modulating activity of canonical Wnt signaling, Notum may also influence canonical Wnt signaling competition with non- canonical Wnt signaling.

## CONCLUSIONS

Our previous studies demonstrated that ablation of Wnt ligand secretion, primarily acting via Wnt/β-catenin signaling, prevents the formation of cartilage giving rise to collapsed and stenotic trachea surrounded by ectopic muscle (Snowball et al., 2015). In the present study, we demonstrated that Wnt5a-mediated non-canonical Wnt signaling supports the timely formation and maturation of the cartilaginous tracheal rings. We also determined functional relationships between Notum and Wnt5a, wherein Notum can inactivate canonical and non-canonical Wnt signaling. Wnt5a deficiency leads to an earlier chondrogenesis, while Notum mutants present a delayed chondrogenesis (Fig 7 c). Furthermore, increased levels of Wnt5a leads to delayed chondrogenesis recapitulating the phenotype observed after deletion of Notum. Thus, non-canonical Wnt signaling is required as a checkpoint to prevent premature mesenchymal condensation, while Notum balances levels of canonical and non-canonical Wnt signaling necessary for the timing of tracheal mesenchymal condensations (Fig 7 d).

### A role for Notum in the regulation of non-canonical Wnt signaling

A large body of literature has focused on the role of Notum as a regulator of Wnt/β-catenin signaling in in vitro studies and different animal models (Kakugawa et al., 2015; Madan et al., 2016; Zhang et al., 2015). Our previous studies have demonstrated a critical role for Notum in regulating canonical Wnt signaling in vitro and in vivo during tracheal development (Gerhardt et al., 2018). Furthermore, since Notum can modulate Wnt/β-catenin, inhibition of Notum might be putative therapeutics for diseases characterized by increased Wnt/β-catenin signaling. As an extracellular molecule, Notum constitutes an excellent druggable target (Bayle et al., 2021).

While it has been demonstrated that Notum downregulates Wnt/β-catenin signaling, less is known of its effect in non-canonical Wnt signaling. Studies in liver fibrosis indicate that in vitro, Notum reduced the Wnt5a-induced pJNK activity in hepatic cells (Li et al., 2019). Our investigations have demonstrated that in vitro Notum prevents the non-canonical response induced by Wnt5a. Our ex-vivo studies support a model whereby increased, Wnt5a-induced-signaling, leads to the activation of non-canonical pathways resulting in a delayed chondrogenic process recapitulating the observed phenotype in the Notum deficient trachea. Thus, it is likely that augmented Wnt5a activity partially accounts for the delayed mesenchymal process in Notum deficient tracheas.

### Wnt5a and Notum influence chondrogenic programs

At the molecular level, we have identified differentially regulated genes when comparing the tracheal prechondrogenic mesenchyme at E11.5 and the chondrogenic mesenchyme at E13.5. Notably, these genes mediate processes related to ECM organization, pathways encoding ECM and ECM-related genes, and cartilage formation. We also determined that several of these ECM-related and chondrogenic genes were precociously expressed at E11.5 in Wnt5a mutants or downregulated at E13.5 after the deletion of Notum such as Osr2. Besides its role in early respiratory tract specification (Rankin et al., 2012), recent studies have demonstrated a role for Osr2 integrating biomechanical signaling sensing the stiffness and ECM changes characteristic of tumors (Zhang et al., 2024). The constituents of the ECM support cell aggregation in chondrogenic nodules by coordinating the availability and diffusion of signals and inhibitors (Ono et al 2009) and restrains cells in a compressed form to sustain differentiation induced by the physical compression (Mammoto et al., 2015). The ECM also facilitates the intracellular communication necessary for condensations by supporting gap junctions between cells and reducing the extracellular space between cells thus, increasing the tissue density (Fowler and Larsson, 2020). Simultaneously, changes in ECM composition will affect the fluidity of the mesenchyme promoting patterning (Huycke et al., 2023; Palmquist et al., 2022). Recent studies have shown that Vangl and Wnt5a are required for the fluidity of the pulmonary mesenchyme during sacculation (Paramore et al., 2024). In the present study, we detected that N-cadherin expression is increased at E11.5 in the prechondrogenic mesenchyme of the Wnt5a mutants. The augmented expression of N-cadherin has been associated with active mesenchymal condensation before overt chondrogenic differentiation, and N-cadherin expression is essential for mesenchymal condensation mediating both cell-cell interactions and formation of intracellular adhesion complex (Delise and Tuan, 2002; Wang et al., 2020). Thus, the increased levels of N-cadherin at E11.5 in Wnt5a deficient trachea further support the concept of Wnt5a as a gatekeeper of the timing of mesenchymal condensations via modulating the expression of N-cadherin, ECM, and chondrogenic related genes.

Furthermore, Notum, attenuates levels of canonical and non-canonical Wnt signaling to balance the activity of both branches of the pathway. In fact, analysis of the gene expression, indicates differential regulation of Wnt signaling after deletion of Notum in the developing trachea, including both Wnt/β catenin dependent and independent targets (Figure 6a). This supports the role of Notum in attenuating both signaling pathways at the transcriptional level, which ultimately will contribute to the proper patterning and timing of the mesenchymal condensations.

### Timing of the condensation and effects on cartilage patterning and shape

After deletion of Wnt5a, the shape and number of cartilages are altered; each ring appears ticker than in the wild type, but the number of rings is reduced. The reduced number of rings might result from geometrical constraints due to the faulty axial elongation observed after the deletion of Wnt5a (Kishimoto et al., 2018; Onesto et al., 2019). Conversely, the shape and timing of the mesenchymal condensation may impact chondrogenesis and the shape of the final cartilage. Studies in zebrafish have demonstrated that the position and longevity of the mesenchymal condensation affect the size and shape of facial cartilages in zebrafish (Paudel et al., 2022). In this model, the interplay of the Fgf and Notch signal is essential to determine the shape of the cartilaginous condensation. In chicken and mouse embryos, Shh mediating signaling is required for proper tracheal ring orientation (Kingsley et al., 2023). Thus, an early condensation may prevent cells from being exposed to signals triggered either by the epithelium or other mesenchymal cells within the trachea or the adjacent heart that will determine each cartilaginous ring’s final shape and size.

On the contrary, Notum is characterized by cartilaginous rings thinner than the wild type, with a mild reduction in the number of cartilaginous rings (Gerhardt et al., 2018). In this model, we demonstrated that mesenchymal condensation is delayed, a phenotype that in the present work is recapitulated ex vivo when control tracheas were treated with Wnt5a conditioned medium (Fig7a). Furthermore, in vivo, reduction of Wnt5a partially rescues the delayed mesenchymal condensation in Notum deficient trachea. Thus, the timing of the mesenchymal condensation has an impact on the shape of the cartilage by altering the exposure of cells to signals required for differentiation and maturation of mesenchymal cells into cartilage.

Finally, it should be taken into consideration the likely heterogeneity of the Sox9 positive cells in the tracheal mesenchyme. The heterogeneity of the cells may affect their response to Wnt5a and Notum signaling during the condensation process; however, this analysis is beyond the scope of this study and will be addressed in a follow up manuscript using single cell sequencing.

### Future directions

While we have demonstrated that mesenchymal Wnt5a partially delays tracheal chondrogenesis by influencing the expression of chondrogenic genes via activation of Camk and JNK, and that an excess of Wnt5a leads to delayed mesenchymal condensation, questions about the molecular mechanism remain unanswered. Specifically, what signaling pathway(s) is/are repressed by Wnt5a to ensure the timely expression of genes required for condensation and cartilaginous differentiation? Is Notum working as a timekeeper by balancing Wnt ligand levels (both canonical and non-canonical) enabling the timely formation of cartilaginous condensations? Do Wnt5a and Notum functionally interact to influence ECM-encoding gene expression to affect the fluidity of the tissue and, thus, the timing of mesenchymal condensations? Will the timing of mesenchymal condensation impact other cell lineages, including the dorsal-ventral patterning and differentiation of the tracheal epithelium? Future studies are guaranteed to answer these questions.

## Supporting information

Table 2 antibodies

Control video

Abc 99 video

JNK inhb video

Control video still image

ABC 99 video still image

KN93 video still image

Jnk inhib video still image

## ACKNOWLEDGEMENTS

We appreciate the comments on the manuscript and the project by Dr. Whitsett and Dr. Chen. The authors would like to thank Chuck Crimmel for assistance with graphic design and Joe Kitzmiller for assistance with video processing. This work was partially supported by NIH-NHLBI R01 144774 and HL156860 to DS.

