## Supplementary material for "Wnt5a and Notum Influence the Temporal Dynamics of Cartilaginous Mesenchymal Condensations in Developing Trachea"

**Table1:**

Primers utilized for genotyping.

**Table 2:**

Antibodies used for the studies.

**Supplementary Figure1:**

Dot plots and density plots display the gating strategy utilized to isolate epithelial (APC+), muscle cells (GFP+), and chondroblasts (APC-GFP-, double negative DN) from the total pool of live cells. Gene expression profile as determined by qRT-PCR confirmed the identity of the different cell populations. Sox9 is highly expressed by APC-GFP- cells. Myocd transcripts are present in GFP+ cells but not detected in APC+ or APC-GFP- cells. Nkx2.1 expression was only detected in APC+ cells.

**Supplementary Figure2:**

RNA scope in situ hybridization demonstrates that levels of Wnt5a are not affected after deletion of Notum. Reciprocally, levels of Notum are not affected after deletion of Wnt5a. Representative images are shown.

**Video:**

Time-lapse imaging depicts the cartilaginous mesenchymal condensation in *Sox9KI eGFP* trachea lung tissue subjected to different conditions. Tissue was isolated at E12.5 and cultured at the air-liquid interphase. Timelapse videos were recorded overnight, 10x images were acquired every 30 minutes, between 24 and 42 hours from the beginning of the incubation. In vehicle treated (DMSO) samples, condensations were observed around 30 hours post incubation (video 1), while Treatment with ABC99, a Notum inhibitor, prevented cartilaginous mesenchymal condensations at 42 hours compared to control (arrow in the video 2). On the other hand, KN93 (calmodulin kinase (CamK) inhibitor) and the JNK inhibitor treatments cause earlier condensation at 24 hours of incubation compared to controls (arrows in videos 3 and 4). Note that neither treatment caused toxicity as pulmonary branching occurred uneventfully (asterisk in the lung) regardless of the chemical addition.
