## Supplementary figures and images for "Wnt5a and Notum Influence the Temporal Dynamics of Cartilaginous Mesenchymal Condensations in Developing Trachea"

### ABC 99 video still image

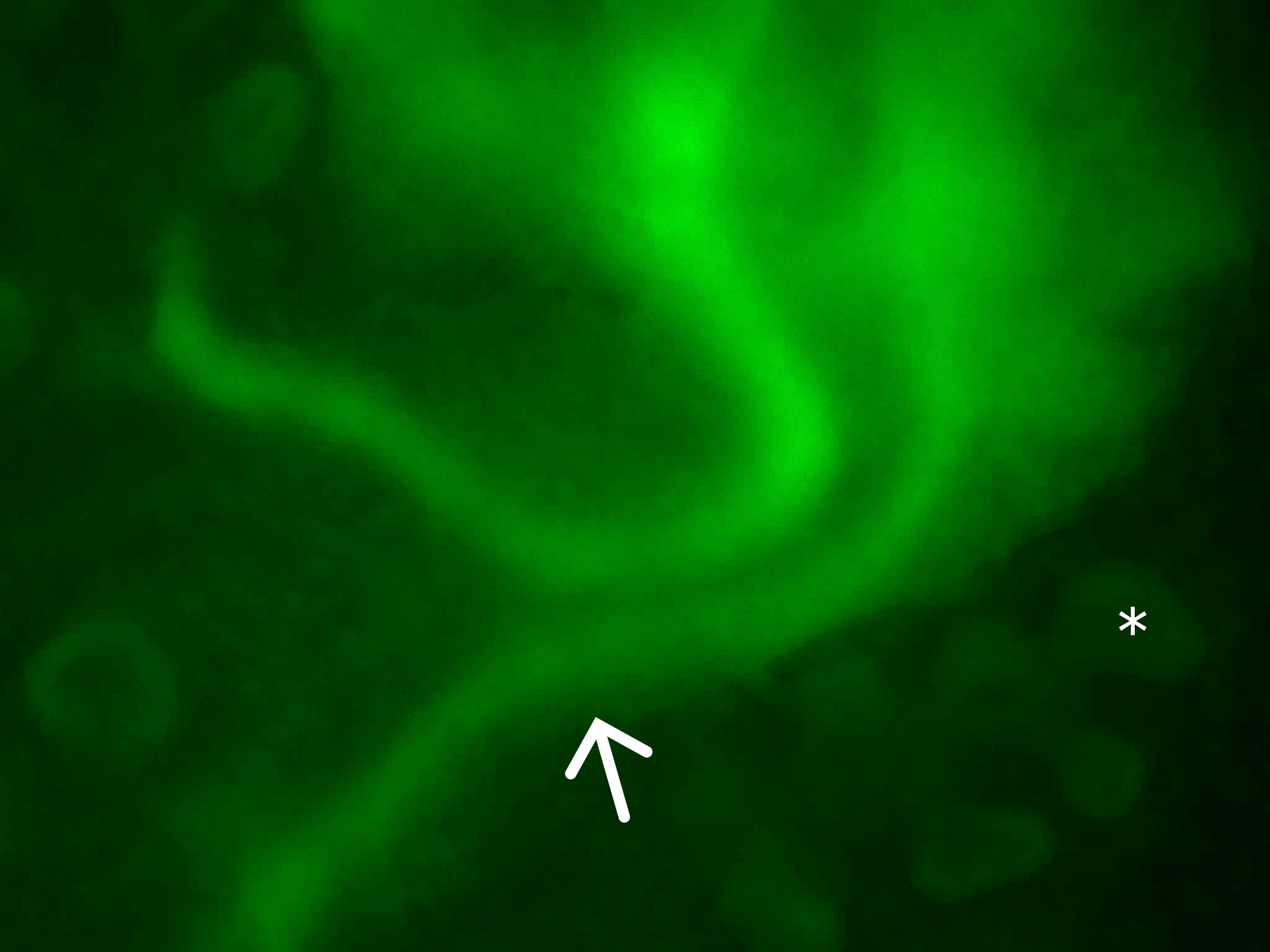

### Control video still image

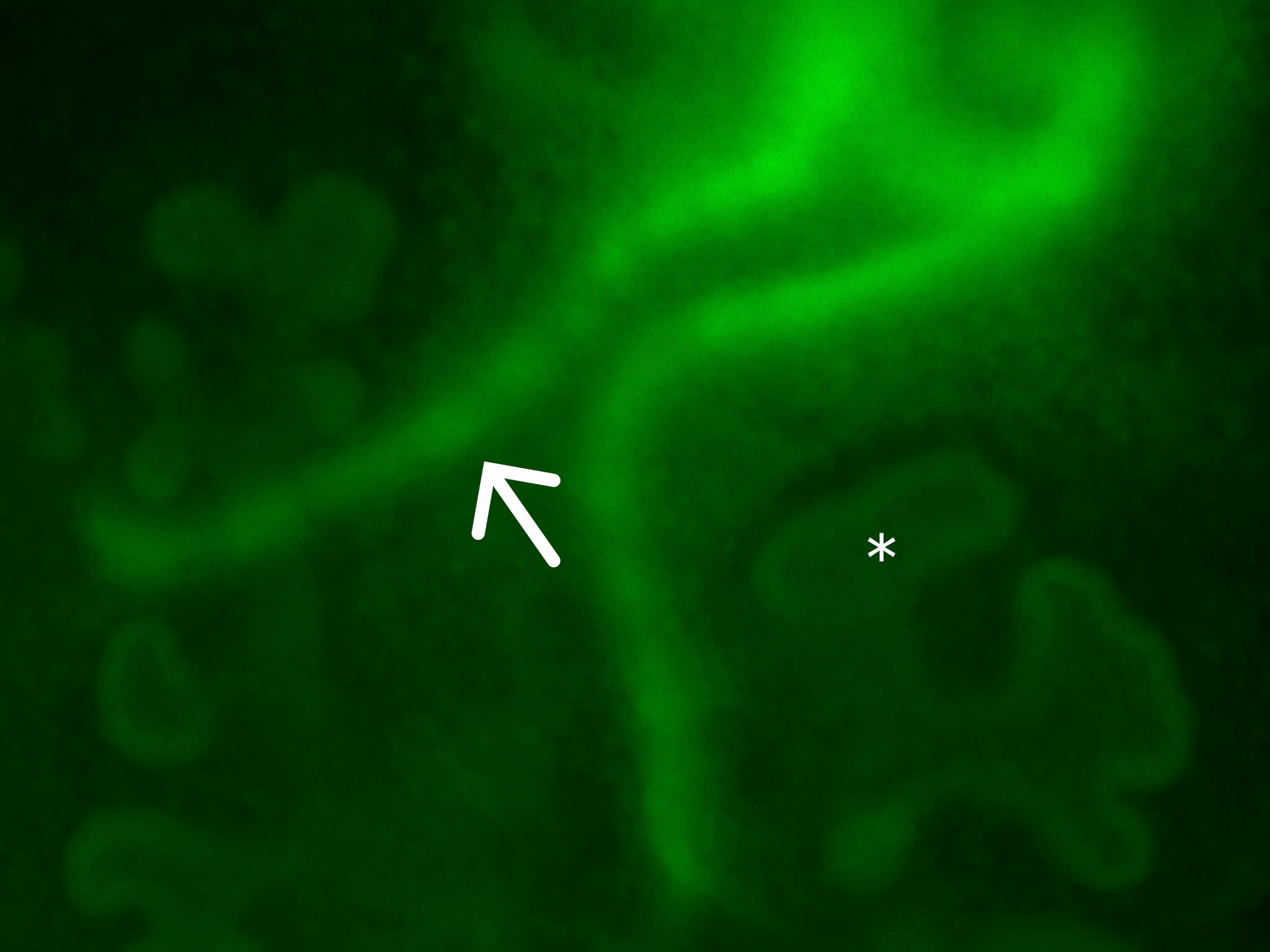

### Jnk inhib video still image

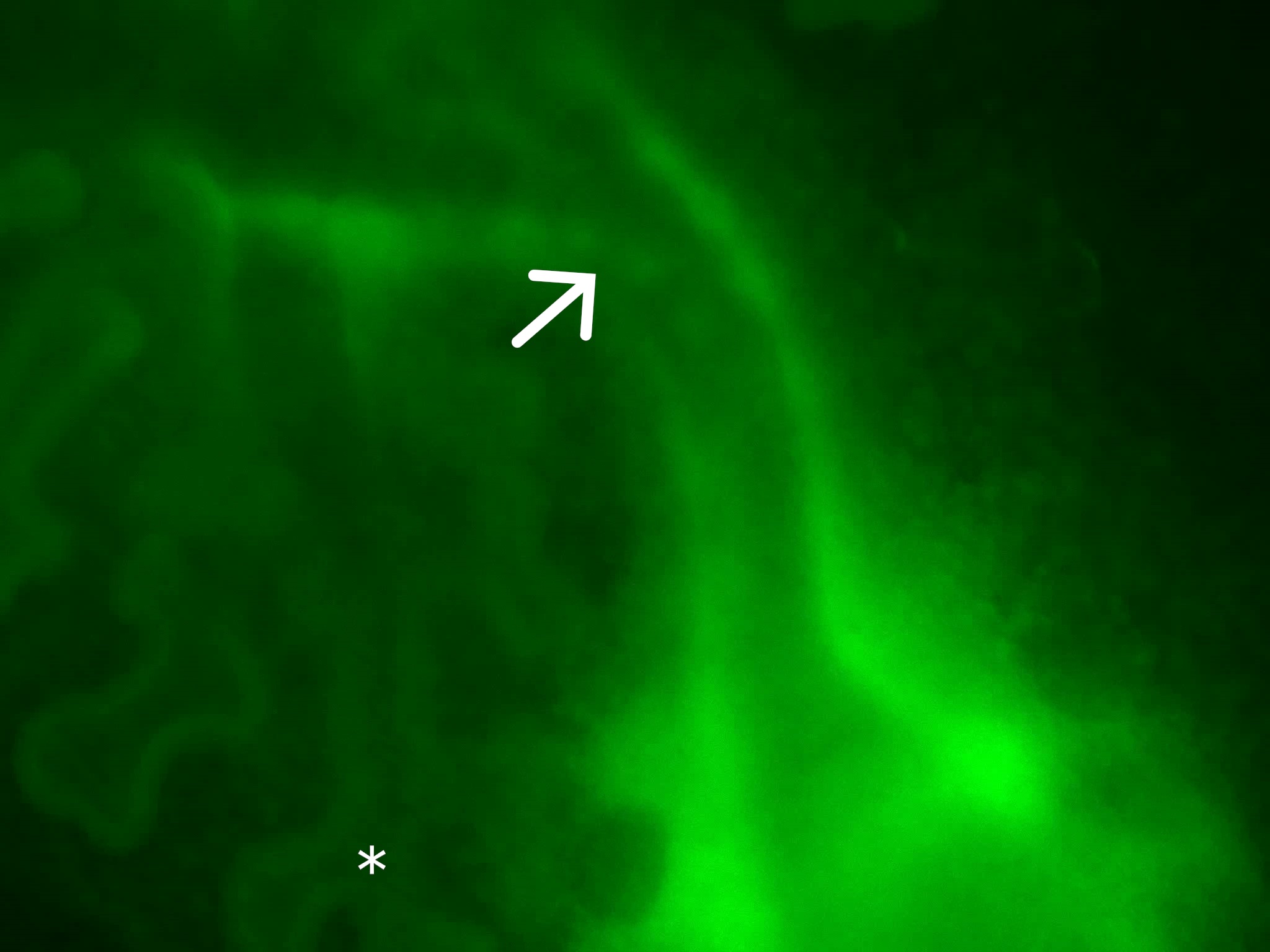

### KN93 video still image

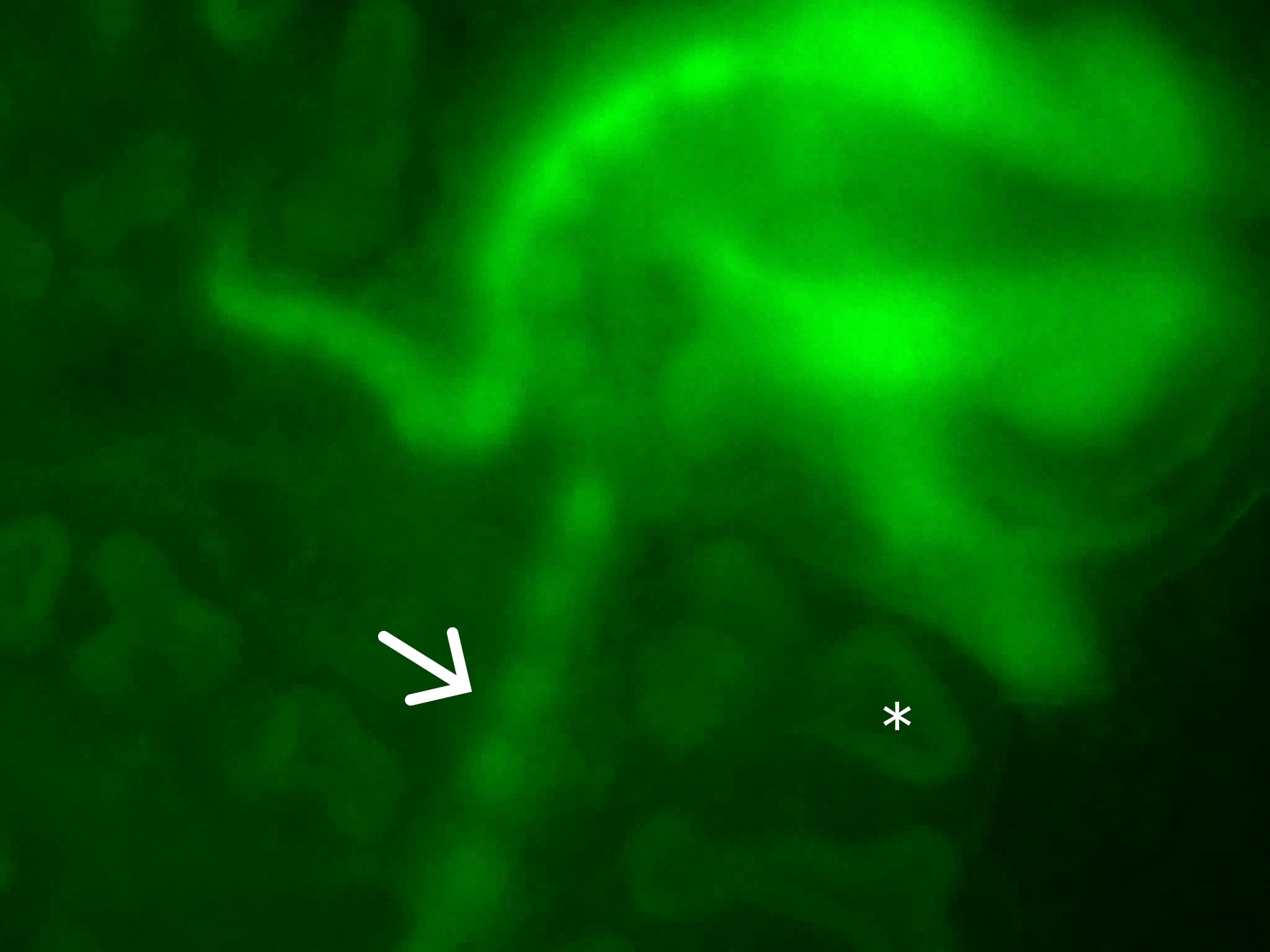

### Supp Fig2

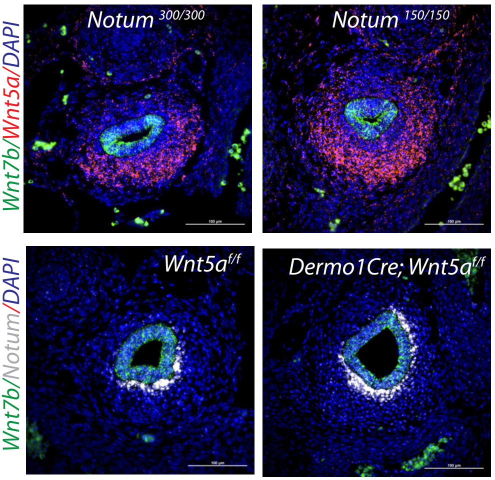

### Supp Figure1

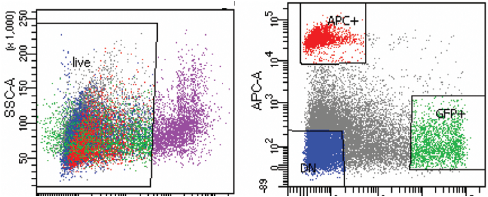


**Expression profile at E13.5 in sorted tracheal cells**

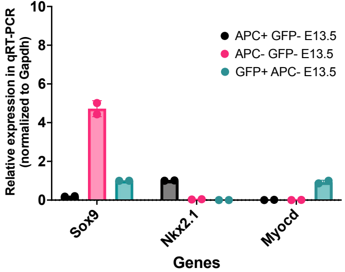
