## Supplementary material for "Wnt5a and Notum Influence the Temporal Dynamics of Cartilaginous Mesenchymal Condensations in Developing Trachea": Table 1

Supplementary Table 1

| | |
|---|---|
| Wnt5af/f | F: GGT GAG GGA CTG GAA GTT GC<br>R: GGA GCA GAT GTT TAT TGC CTT C |
| Dermo1Cre | F: TGC CAC GAC CAA GTG ACA GCA ATG<br>R: AGA GAC GGA AAT CCA TCG CTC G |
| Notum | F: CTG ACA GCA TGC TCT GTG CG<br>R: ACT ATT CTG CAG ACC GAG CCA GTC |
| Sox9KIeGFP | 1: GAG GGG CTT GTC TCC AG<br>2: ACA CCG GCC TTA TTC CAA G<br>3: GGC AGC TAC TCT TGA AAT CCA |
| Gamma SMA | F: CCT ACG GCG TGC AGT GCT TCA GC<br>R: CGG CGA GCT GCA CGC TGC GTC CTC |
